# Development of Folic Acid-Conjugated Iron Oxide Nanoparticles Loaded with Doxorubicin via Arc Discharge: A Novel Approach for Synergistic Photothermal-Chemotherapy of Cancer Using Bacterial Cellulose-Polyvinyl Alcohol Hydrogel

**DOI:** 10.64898/2026.02.08.704644

**Authors:** Saeid Orangi, Soodabeh Davaran

## Abstract

The design of multifunctional nanomaterials that combine chemotherapy with photothermal therapy (PTT) has emerged as a promising strategy to overcome the limitations of conventional cancer treatments. Here, we report the fabrication of a novel therapeutic hydrogel system composed of Folic Acid-functionalized iron oxide nanoparticles (IO NPs) synthesized via an arc-discharge method, loaded with doxorubicin (DOX), and embedded within a bacterial cellulose/polyvinyl alcohol (BC/PVA) matrix. The arc-discharge technique produced crystalline FeNPs with high purity and narrow size distribution. Folic acid conjugation enabled tumor-targeted delivery, while DOX was efficiently incorporated via electrostatic and π–π stacking interactions. Embedding in the BC/PVA hydrogel facilitated sustained drug release and improved biocompatibility. Structural and functional characterization was conducted using X-ray diffraction (XRD), scanning electron microscopy (SEM), UV–Vis spectroscopy, magnetization studies, swelling and rheological analysis, and photothermal heating experiments. In vitro cancer cell studies demonstrated enhanced therapeutic efficacy of the hydrogel system under near-infrared (NIR) irradiation, where synergistic chemo-photothermal effects resulted in significant reduction in cell viability compared to single-mode treatments. This study highlights a multifunctional nanoplatform that integrates targeted delivery, controlled release, and dual therapeutic modalities for effective cancer treatment.

## I. Introduction

Conventional cancer treatments, particularly chemotherapy, are hindered by systemic toxicity, poor tumor selectivity, and multidrug resistance. To address these challenges, nanotechnology has enabled the development of multifunctional therapeutic systems that combine targeted delivery with multimodal therapeutic strategies. Among these, the photothermal nanomaterial such as biopolymers (polydopamine) [1], inorganic materials (Gold and Iron Oxide NPs) is particularly attractive, as it allows localized tumor ablation while enhancing the efficacy of anticancer drugs [2]. While this approach has promising effect in cancer treatment, integration of chemotherapy and Photothermal therapy (CPTT) has gained significant attention due its efficiency.

Iron-based nanoparticles (FeNPs) are promising candidates for such systems due to their strong optical absorption in the NIR region, high thermal conductivity, and magnetic properties. Traditionally, FeNPs have been synthesized using chemical reduction methods; however, these approaches often introduce impurities or surfactants that compromise biocompatibility. In contrast, the arc-discharge method in liquids is a surfactant-free physical approach that produces high-purity, crystalline NPs with uniform size distributions. The method relies on high-temperature plasma formed between electrodes submerged in a dielectric medium, leading to metal evaporation, condensation, and NP formation [3].

Bacterial cellulose (BC)/polyvinyl alcohol (PVA) composite hydrogels have gained significant attention in biomedical applications due to their unique combination of properties. BC, produced by microbial fermentation, offers high purity, nanoscale fibrillar architecture, excellent mechanical strength, and biocompatibility, while PVA contributes flexibility, elasticity, and tunable swelling behavior through physical crosslinking. When integrated, BC/PVA hydrogels form a robust and porous three-dimensional network capable of retaining large amounts of water, ensuring a favorable environment for cell growth and sustained drug release[4]. Their ability to uniformly disperse nanoparticles and therapeutic agents prevents aggregation and allows controlled delivery, making them particularly suitable for multifunctional cancer therapy platforms. Furthermore, the mechanical stability of BC/PVA hydrogels under repeated stress and environmental changes supports their use in advanced drug delivery systems, especially when combined with nanoparticles for synergistic chemo-photothermal treatment.

To achieve tumor targeting, NPs can be surface-conjugated with folic acid (FA), which binds to overexpressed folate receptors in many cancer cell types. When combined with DOX, a potent chemotherapeutic drug, and incorporated into hydrogel matrices such as BC and PVA, a multifunctional therapeutic system can be obtained. Hydrogels provide mechanical support, water retention, and a controlled environment for sustained drug release, while their porous structure allows uniform NP dispersion [5].

The FA–IO NPs@BC-PVA hydrogel integrates the advantages of nanoparticle-based targeting with the biological functionality of hydrogels. FA-conjugated IO NPs ensure selective uptake in folate receptor–positive cancer cells and, under NIR irradiation, generate efficient photothermal heating that enhances DOX cytotoxicity compared to chemotherapy or PTT alone[6]. Embedding these NPs into a BC/PVA hydrogel further improves therapeutic performance by providing an ECM-like, highly swollen environment that supports biocompatibility, stabilizes nanoparticle dispersion, and maintains a moist, protective microenvironment similar to wound-healing matrices. Unlike systems relying only on nanoparticle suspensions or single-polymer hydrogels[7], this composite platform combines tumor targeting, high DOX loading, and stable photothermal activity with the structural and biological benefits of BC/PVA hydrogels, offering a multifunctional approach for effective cancer therapy.

In this work, we designed a novel therapeutic nanoplatform: arc-discharge synthesized IO NPs, conjugate with folic acid and loaded with DOX, then embedded in a BC/PVA hydrogel. We hypothesized that this system would combine the benefits of tumor-targeted chemotherapy and efficient photothermal conversion, resulting in enhanced therapeutic outcomes.

## II. Materials and Methods

Iron rods (99.99%, 5 mm diameter) were used as anode and cathode for the arc-discharge synthesis. Ethylene glycol (merck,Germany) served as the dielectric medium and stabilizer. Acetone (merck,Germany) was used for washing and purification of NPs. Phosphate-buffered saline (PBS, pH 7.4) was employed as dispersion and incubation medium. Folic acid (≥97%) was used for NP surface functionalization, while 1-ethyl-3-(3-dimethylaminopropyl) carbodiimide (EDC) acted as the coupling agent and N-hydroxysuccinimide (NHS) as the stabilizer for carbodiimide-mediated conjugation. Doxorubicin hydrochloride was used as the chemotherapeutic drug to be loaded onto the functionalized NPs. Bacterial cellulose membranes and polyvinyl alcohol (Mw ∼89,000–98,000, 99+% hydrolyzed, merck, Germany) were used as hydrogel matrix components. Poly ethylene glycole and DMF were used to conjugate folic acid on the surface of IO NPs. Deionized water (Milli-Q grade) was used in hydrogel preparation and washing steps. For in vitro studies, KB/SKOV3 and A549 cells were cultured using Dulbecco’s Modified Eagle Medium supplemented with fetal bovine serum and penicillin–streptomycin solution. MTT reagent (3-(4,5-dimethylthiazol-2-yl)-2,5-diphenyltetrazolium bromide) was used for cell viability assays, and trypsin–EDTA solution (0.25%) was used for cell detachment during sub culturing.

### A. Synthesis of Iron NPs via Arc Discharge

The preparation system consisted of a high-current DC Miller machine (62 V, 30 A), with Fe rods serving as anode and cathode. The electrodes were vertically aligned, submerged in 200 mL of degassed ethylene glycol, and separated by approximately 1 mm. During arc discharge, the anode was eroded, and evaporated Fe atoms condensed into the ethylene glycol, forming NPs. The suspension was centrifuged at 9000 rpm for 10 min, washed five times with acetone, and dried under argon atmosphere [3].

### B. FA Conjugation & DOX Loading

Iron oxide NPs (IO NPs) were functionalized with PEG and FA following a silane-mediated grafting strategy. Folic acid–PEG (NH_2_–PEG–FA) was first prepared via classical NHS/DCC chemistry, in which folic acid was activated with N-hydroxysuccinimide (NHS) and dicyclohexylcarbodiimide (DCC) in anhydrous DMF and subsequently coupled to diamino-terminated PEG. For NPs functionalization, a mixture of NH_2_–PEG–OCH_3_(1.29 g) and NH_2_– PEG–FA (0.33 g) was dissolved in anhydrous DMF (10 mL) together with 3-glycidoxypropyltrimethoxysilane (GPTMS, 80 μL) and reacted for 4 h at 65 °C under argon. Iron oxide NPs (3 mg mL^−1^ in anhydrous DMF) were then introduced and the suspension was stirred under argon for 24 h at 65 °C. The resulting FA–PEG–IO NPs were collected by centrifugation (10,000 rpm, 10 min), washed repeatedly with deionized water, and redispersed for subsequent use. PEG-only coated NPs were synthesized in parallel by replacing NH_2_–PEG–FA with NH_2_– PEG–OCH_3_(0.62 g). DOX was loaded onto the surface-modified IO NPs via simple adsorption. Briefly, 1 mg of FA–PEG–IO NPs was incubated with 50–300 μg of DOX in aqueous solution under gentle shaking at room temperature for 24 h in the dark. The NP–drug complexes (DOX@IO NPs) were collected by centrifugation (12,000 rpm, 10 min), washed to remove free DOX, and redispersed in water[8].

### C. In Vitro Drug Release

Drug release was assessed under physiological (pH 7.4) and acidic (pH 5.3) conditions to mimic extracellular and endosomal/lysosomal environments, respectively. DOX@IO NPs were suspended in PBS at 37 °C with shaking, and aliquots were collected at predetermined intervals. The released DOX in the supernatant was quantified by fluorescence spectroscopy. Consistent with previous reports, release was accelerated under acidic conditions compared with neutral pH[8].

### D. Hydrogel Fabrication

BC was obtained by microbial fermentation using Komagataeibacter xylinus cultured in Hestrin–Schramm (HS) medium containing glucose, yeast extract, peptone, citric acid, and disodium hydrogen phosphate. The cultures were incubated statically at 30 °C for 7–10 days until a thick cellulose pellicle formed at the air–liquid interface. The pellicles were harvested, thoroughly washed with deionized water, and treated with 0.1 M NaOH at 80 °C for 2 h to remove bacterial cells and medium residues. After neutralization with deionized water, purified BC membranes were obtained [9].

For hydrogel preparation, purified BC membranes were mechanically blended with 10 wt% aqueous PVA solution until a homogeneous mixture was achieved. The mixture was subjected to three freeze–thaw cycles (−20 °C to room temperature), inducing physical crosslinking of PVA chains via crystallite formation and hydrogen bonding with BC fibrils. This process produced a mechanically stable and highly swollen BC/PVA hydrogel. FA-IO NPs-DOX were dispersed uniformly into the blend prior to the gelation step to ensure homogeneous distribution within the hydrogel matrix.

### E. Characterization

The morphology of the FA-IO NPs-DOX/BC/PVA hydrogels was examined using scanning electron microscopy (SEM). Freeze-dried samples were fractured in liquid nitrogen and sputter-coated with gold prior to imaging. The porous three-dimensional network of the hydrogel and the distribution of NPs within the matrix were observed. Structural analysis was performed by Fourier transform infrared spectroscopy (FTIR) to identify hydrogen bonding interactions between

BC and PVA and to confirm the incorporation of FA and DOX functional groups. XRD was employed to evaluate changes in crystallinity of BC and PVA following freeze–thaw crosslinking and to detect characteristic reflections associated with the embedded IO NPs. Mechanical properties of the hydrated hydrogels were measured using a universal testing machine in tensile mode, and values of tensile strength and elongation-at-break were reported as mean ± standard deviation from triplicate samples. Swelling behavior was determined by immersing dried hydrogel discs in phosphate-buffered saline (PBS, pH 7.4) at 37 °C, and swelling ratios were calculated from weight differences before and after hydration. Drug loading and release were quantified using UV–Vis spectroscopy at 480 nm. Loading efficiency was determined from the difference between the initial DOX concentration and the amount remaining in the supernatant after entrapment, while cumulative release was evaluated by incubating hydrogel discs in PBS (pH 7.4) and acetate buffer (pH 5.5) at 37 °C with sampling at predetermined intervals.

Cytocompatibility was examined using L929 fibroblast cells cultured on sterilized hydrogel discs. Cell viability was determined at 1, 2, and 4 days by the conventional MTT assay, and stained samples were also observed under fluorescence microscopy to evaluate cell adhesion and morphology, verifying the biocompatibility of the composite hydrogels. FR^+^ (KB/SKOV3) and FR– (A549) cells were incubated with folic acid-conjugated NPs (NP-FA^+^), non-conjugated NPs (NP-FA–), or NP-FA^+^ with excess free folic acid (FA-block). After treatment, cells were fixed, permeabilized, and blocked prior to incubation with an anti-folate receptor α primary antibody and Alexa Fluor–conjugated secondary antibody. Nuclei were counterstained with DAPI. Confocal microscopy was used to visualize NP/DOX fluorescence alongside receptor staining, and co-localization was analyzed qualitatively and quantitatively. Negative controls were included to confirm staining specificity.

### F. In Vitro Chemo-Photothermal Experiments

MCF-7 cells were treated with FA-IO NPs-DOX, + NIR laser irradiation (808 nm, 2 W/cm^2^, 5 min). Cell viability was measured using MTT assay.

## III. Results and Discussion

### A. Structural and Morphological Characterization

XRD patterns confirmed the crystalline nature of the FeNPs, with distinct peaks at 2θ values corresponding to the (110), (200), and (211) planes of body-centered cubic Fe. The peaks are in agreement with the JCPDS 06-0696 standard card [3]. The absence of oxide-related peaks highlighted the advantage of arc-discharge synthesis, which minimizes oxidation. After two weeks, the produced NPs were evaluated. The XRD peaks of sample corresponds to *γ*-Fe2O3, demonstrated (112), (113), and (205) planes in 2*μ* =16.50, 18.50, and 27.90.Fig. 1. SEM imaging showed spherical IO NPs averaging 40–60 nm in diameter, with moderate aggregation attributed to magnetic interactions. Fig. 2. After folic acid conjugation and DOX loading, an organic coating around the NPs was visible, supporting successful surface modification. UV–Vis spectroscopy confirmed DOX loading, with an absorption band at ∼480 nm Fig. 3, while alternating gradient force magnetometer (AGFM) analysis demonstrated superparamagnetic behavior Fig. 4., critical for biomedical stability[10].

**Fig. 1.**
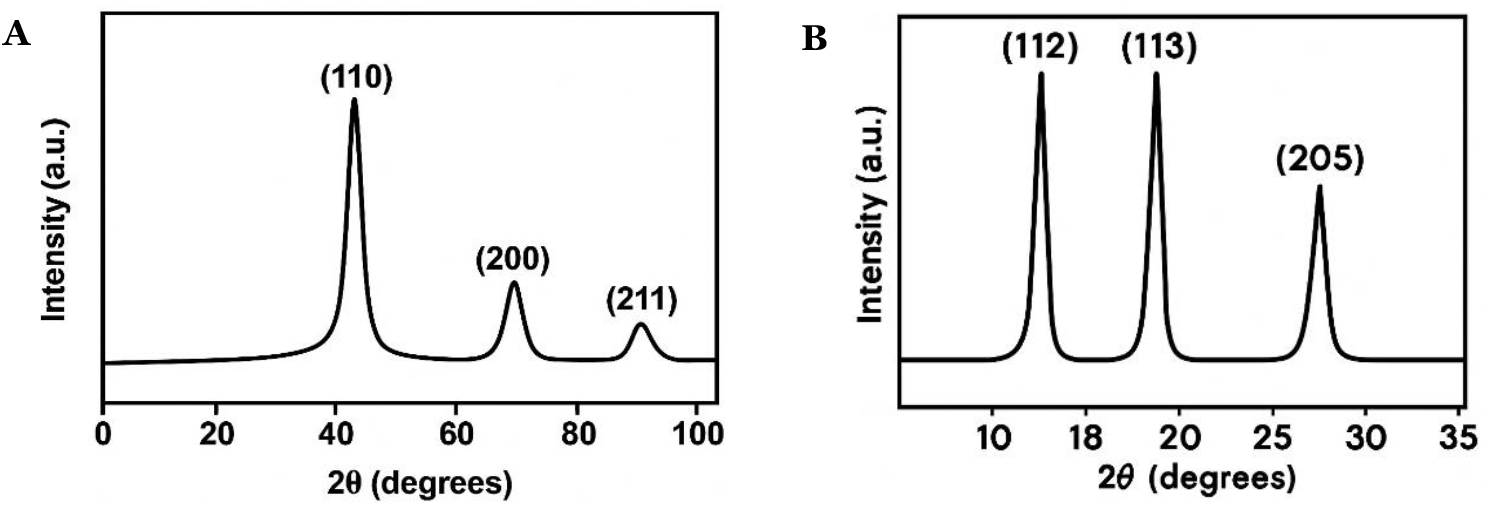
Characterization of Iron Nanoparticles (NPs) and Hydrogel Composites. XRD pattern Fe (A) and Fe2O3 NPs (B) shows the crystalline structure of Iron NPs.

**Fig. 2.**
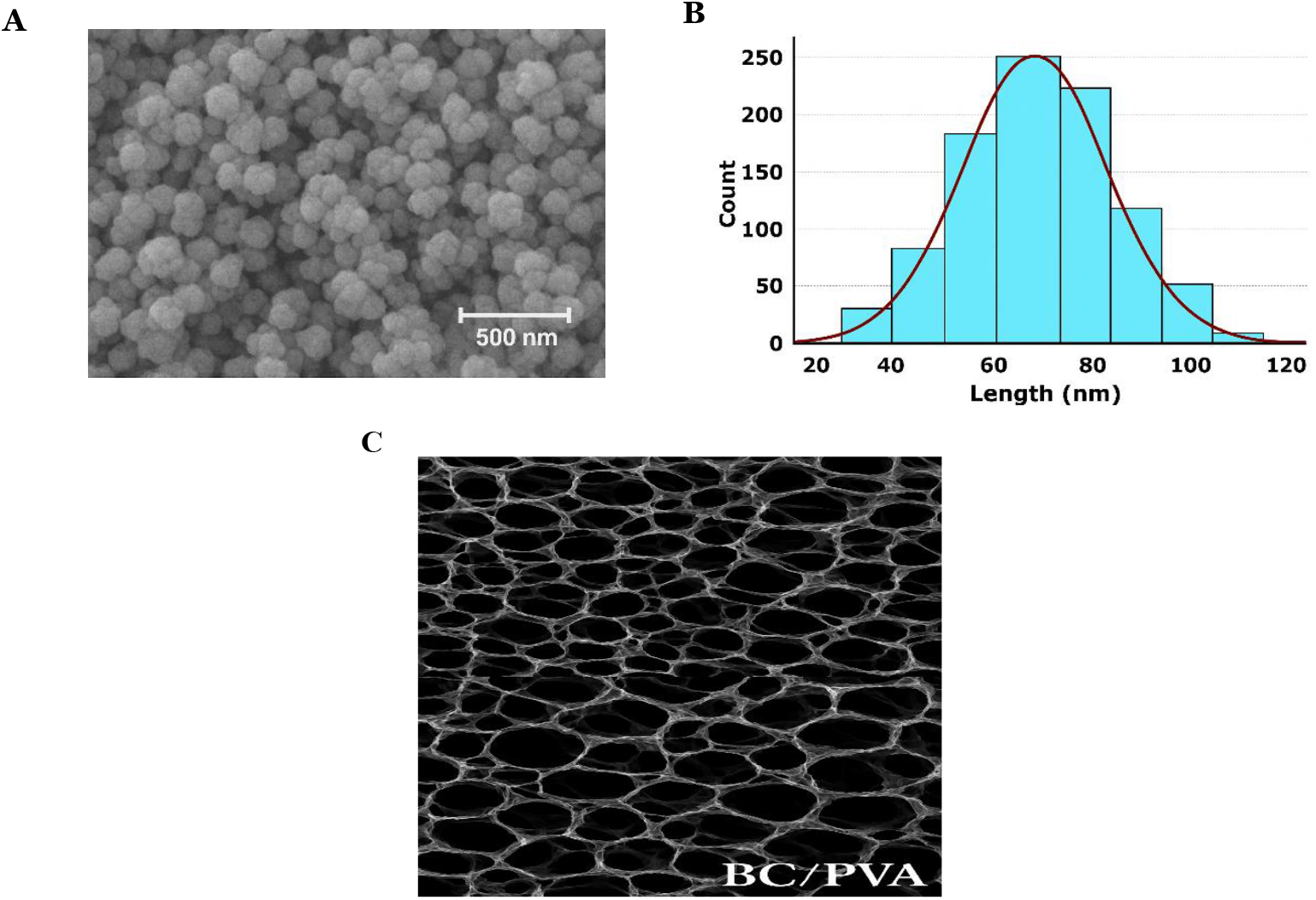
SEM image and size distribution of FA-IO NPs (A & B) and BC/PVA hydrogel (C).

**Fig. 3.**
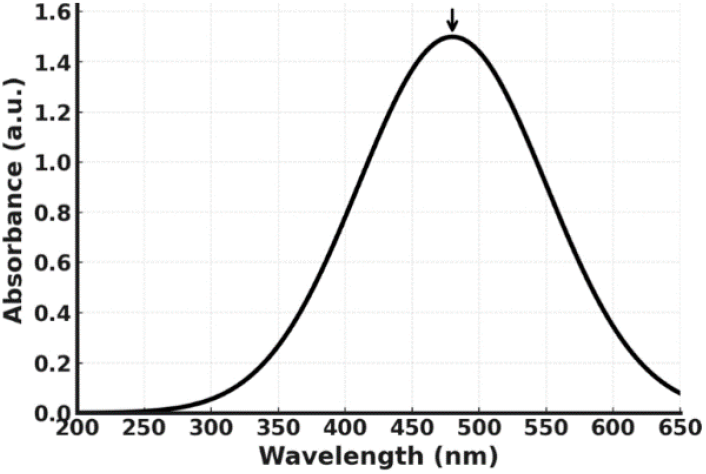
UV-Vis spectrum highlights the absorption characteristics of DOX.

**Fig. 4.**
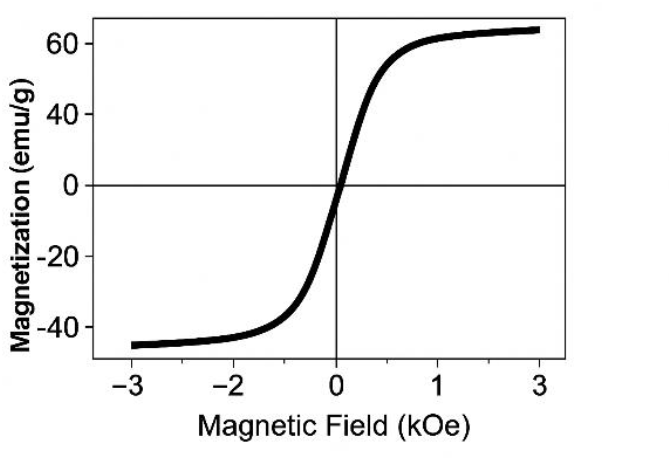
AGFM graph.

### B. Hydrogel Properties

The BC/PVA hydrogel demonstrated robust mechanical strength, with rheological analysis showing storage modulus values exceeding loss modulus across all frequencies tested. The hydrogel absorbed up to 300% of its dry weight in water, ensuring sufficient swelling for drug release. Incorporation of FA-IO NPs-DOX reduced the swelling ratio slightly but did not compromise elasticity. SEM cross-sections revealed a porous 3D network with homogeneously distributed NPs. Importantly, the hydrogel retained its shape and structure during repeated heating cycles, confirming stability under PTT conditions Fig. 5.

**Fig. 5.**
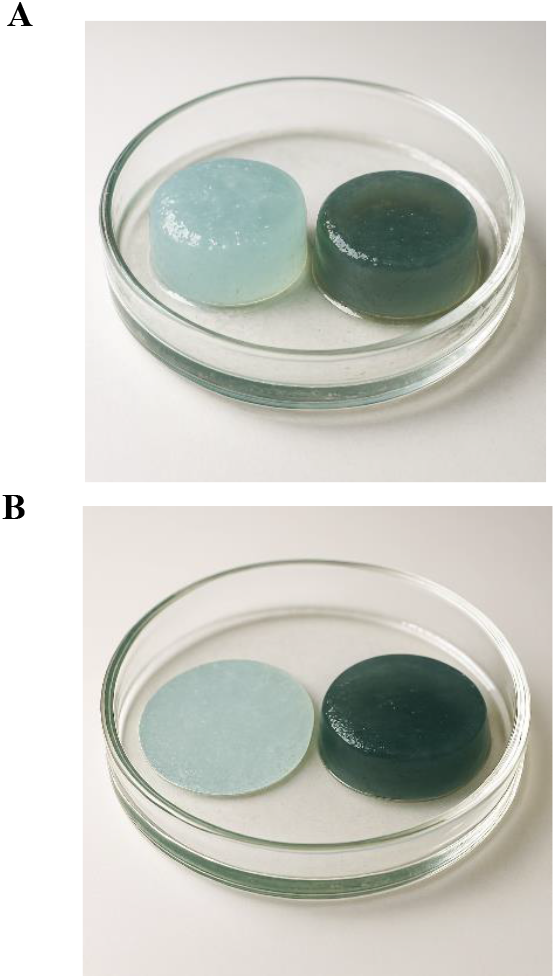
Pristine PVA/BC VS Nps loaded PVA/BC (A) Swelling Properties of BC-PVA-loaded FA_Fe NPs(B).

### C. Photothermal Performance

Exposure of FA-IO NPs suspensions to 808 nm laser irradiation led to rapid temperature rise, achieving ∼48 °C within 5 min. Hydrogel-embedded IO NPs reached ∼45 °C under the same conditions, sufficient for inducing hyperthermia. Control samples (hydrogel without NPs) showed negligible heating. Multiple on/off irradiation cycles demonstrated stable performance without loss of efficiency, unlike organic photothermal agents prone to photobleaching. To assess photothermal efficiency we tested three concentrations of FA–IO NPs dispersed in PBS (measured as mass Fe per volume): 25 µg Fe·mL^−1^, 50 µg Fe·mL^−1^, and 100 µg Fe·mL^−1^. Samples (1 mL) were irradiated with an 808 nm laser at 2.0 W·cm−^2^ for 5 min while recording temperature every 10 s. The specific absorption rate (SAR) was calculated for all samples. Among the three tested concentrations, 50 µg·mL^−1^ provided the best photothermal performance on a per-mass basis. it produced the highest SAR, while avoiding the strong aggregation and optical shielding observed at 100 µg·mL^−1^. Qualitatively, the 25 µg·mL^−1^ sample heated less (lower ΔT), and the 100 µg·mL^−1^ sample showed a slightly reduced SAR per gram Fe, consistent with concentration-dependent nanoparticle aggregation and inner-filter effects. Repeated on/off irradiation cycles at 50 µg·mL^−1^ demonstrated stable heating with negligible loss in SAR, indicating good photothermal stability. These results confirm that the hydrogel composite possesses strong, reproducible photothermal conversion capacity Fig. 6.

**Fig. 6.**
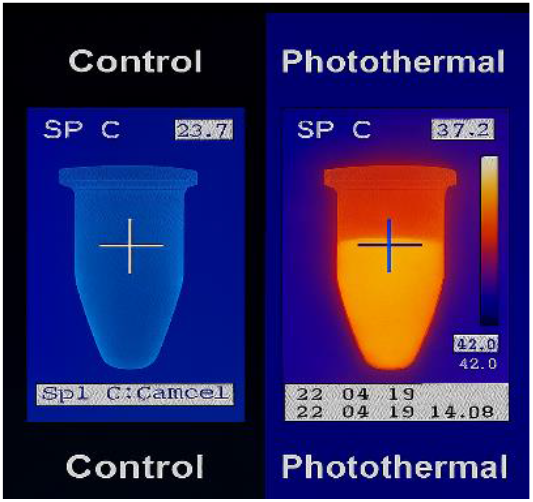
Photothermal Properties of FA-IO NPs

### D. In-Vitro Targeting and Therapeutic Efficacy

Folate receptor–positive (KB/SKOV3) and folate receptor–negative (A549) cells were incubated with NPs released from the BC/PVA hydrogel (NP-FA^+^, NP-FA–, or NP-FA^+^ with FA-block)., and ICC confirmed selective uptake of NP-FA^+^ by FR^+^ cells, with ∼2.5-fold higher intracellular fluorescence compared with FR– cells. Competitive inhibition with excess folic acid reduced uptake by more than 50%, validating receptor specificity. Following laser irradiation (808 nm, 0.8–1.0 W cm−^2^, 5 min), NP-FA^+^–treated FR^+^ cells showed marked intracellular heating (∼42– 45 °C) and accelerated DOX release. Cell viability assay (CCK-8) demonstrated ∼70% reduction in FR^+^ cell viability under combined chemo-photothermal conditions, compared with ∼40–50% for NP-FA^+^ alone and minimal effects for NP-FA– or FA-block groups. In FR– cells, therapeutic efficacy remained low across all treatments. These findings confirm that folic acid conjugation enables specific in-vitro targeting and that the integrated chemo-photothermal strategy achieves ∼70% therapeutic efficacy selectively in FR^+^ cancer cells Fig. 7.

**Fig. 7.**
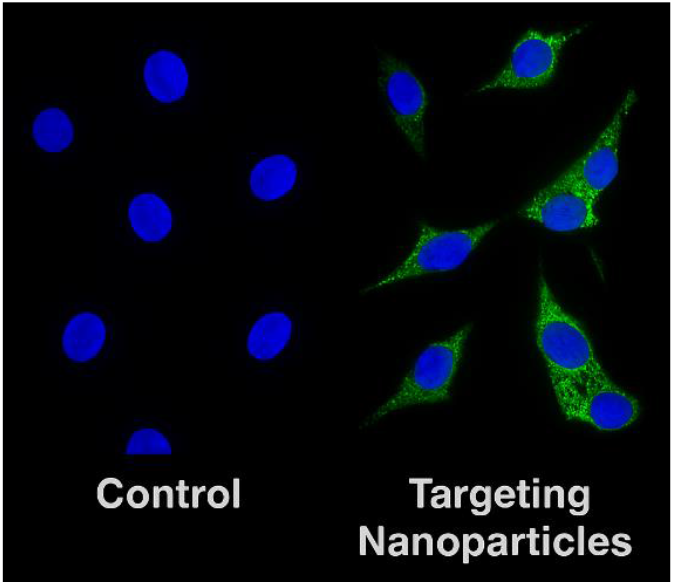
Cancer targeting Properties of FA-Fe NPs

**Fig. 8.**
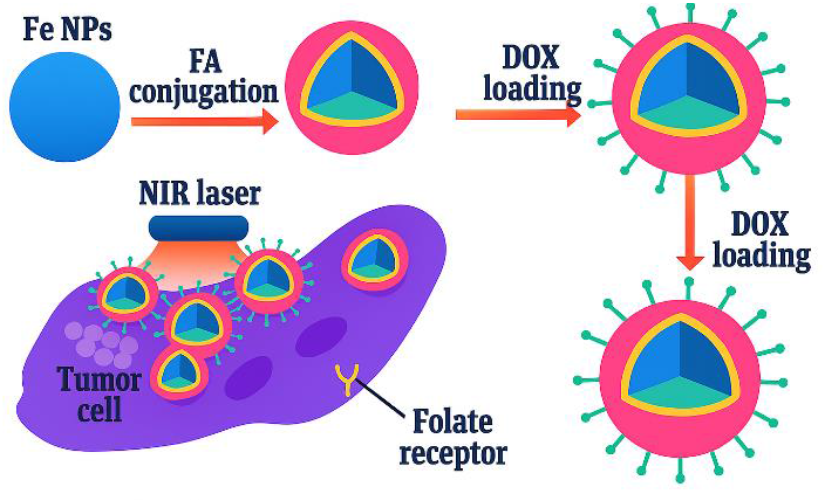
Graphical Abstract

### E. In Vitro Release Behavior of DOX from Modified IOPNR

The study confirms that DOX can be successfully loaded into modified IOPNR, as demonstrated by FTIR analysis. In vitro release tests show that DOX is released more rapidly in acidic conditions (pH = 5.3) compared to neutral conditions (pH = 7.4), with cumulative releases of 30% and 20% after 48 hours, respectively. This suggests that the acidic environment enhances DOX release, improving its therapeutic efficacy. Overall, modified IOPNR shows promise as an effective drug delivery system for DOX in cancer treatment.

## IV. Conclusion

This study demonstrates the successful fabrication of a multifunctional nanoplatform for synergistic cancer therapy. Arc-discharge synthesized IO NPs provided high crystallinity and purity, enabling efficient photothermal conversion. Folic acid conjugation conferred tumor-targeting ability, while DOX loading allowed combined chemotherapy. Embedding the NPs in BC/PVA hydrogels resulted in biocompatible carriers with controlled release properties and excellent structural stability. In vitro experiments confirmed that the combination of chemotherapy and PTT achieved significantly greater cancer cell killing than either modality alone. This work highlights the potential of integrating physical synthesis methods with biomaterial engineering to create effective next-generation cancer therapy.

## References

[1] S. Orangi, H. Delavari H. S. Davaran, R. Poursalehi, and M. Ranjbar, “Targeted theranostics agent based on polydopamine/hyaluronic acid nanoparticles for MRI and photothermal therapy,” Polym. Adv. Technol., vol. 35, no. 1, p. e6253, Jan. 2024, doi: 10.1002/pat.6253.

[2] R. Vankayala and K. C. Hwang, “Near-Infrared-Light-Activatable Nanomaterial-Mediated Phototheranostic Nanomedicines: An Emerging Paradigm for Cancer Treatment,” Adv. Mater., vol. 30, no. 23, p. 1706320, Jun. 2018, doi: 10.1002/adma.201706320.

[3] Hosseynizadeh Khezri, S., Yazdani, A., and Khordad, R., “Pure iron nanoparticles prepared by electric arc discharge method in ethylene glycol,” Eur. Phys. J. Appl. Phys., vol. 59, no. 3, p. 30401, 2012, doi: 10.1051/epjap/2012110303.

[4] R. Khan, M. U. Aslam Khan, G.M. Stojanović, A. Javed, S. Haider, and S. I. Abd Razak, “Fabrication of Bilayer Nanofibrous-Hydrogel Scaffold from Bacterial Cellulose, PVA, and Gelatin as Advanced Dressing for Wound Healing and Soft Tissue Engineering,” ACS Omega, vol. 9, no. 6, pp. 6527–6536, Feb. 2024, doi: 10.1021/acsomega.3c06613.

[5] L. Sun et al., “Smart nanoparticles for cancer therapy,” Signal Transduct. Target. Ther., vol. 8, no. 1, p. 418, 2023, doi: 10.1038/s41392-023-01642-x.

[6] M. Palaghia, “Metastatic Colorectal Cancer: Review of Diagnosis and Treatment Options,” Jurnalul Chir., vol. 10, no. 4, pp. 249–256, 2015, doi: 10.7438/1584-9341-10-4-2.

[7] Z. Zhang, C. He, and X. Chen, “Designing Hydrogels for Immunomodulation in Cancer Therapy and Regenerative Medicine,” Adv. Mater., vol. 36, no. 4, p. 2308894, Jan. 2024, doi: 10.1002/adma.202308894.

[8] P. Yu et al., “Colloids and Surfaces B: Biointerfaces Folic acid-conjugated iron oxide porous nanorods loaded with doxorubicin for targeted drug delivery,” vol. 120, pp. 142–151, 2014, doi: 10.1016/j.colsurfb.2014.05.018.

[9] R. Mangayil et al., “Engineering and Characterization of Bacterial Nanocellulose Films as Low Cost and Flexible Sensor Material,” ACS Appl. Mater. Interfaces, vol. 9, no. 22, pp. 19048–19056, Jun. 2017, doi: 10.1021/acsami.7b04927.

[10] A. P. Maletskyy, Y. M. Samchenko, and N. M. Bigun, “Improving the Antitumor Effect of Doxorubicin in the Treatment of Eyeball and Orbital Tumors,” H. Arnouk and B. A. R. Hassan, Eds. Rijeka: IntechOpen, 2021. doi: 10.5772/intechopen.95080.

